# Optical Control of Cytokine Signaling via Bioinspired, Polymer-Induced Latency

**DOI:** 10.1101/2020.02.13.948240

**Authors:** Lacey A Perdue, Priscilla Do, Camille David, Andrew Chyong, Anna Kellner, Amanda Ruggieri, Hye Ryong Kim, Khalid Salaita, Gregory B Lesinski, Christopher C Porter, Erik C Dreaden

## Abstract

Cytokine signaling is challenging to study and therapeutically exploit as the effects of these protein are often pleiotropic. A subset of cytokines can, however, exert signal specificity via association with latency-inducing proteins which cage the cytokine until disrupted by discreet biological stimuli. Inspired by this precision, here we describe a strategy for synthetic induction of cytokine latency via modification with photo-labile polymers that mimic latency while attached, then restore protein activity in response to light, thus controlling the magnitude, duration, and location of cytokine signals. We characterize the high dynamic range of latent cytokine activity modulation and find that polymer-induced latency, alone, can prolong *in vivo* circulation and bias receptor subunit binding. We further show that protein de-repression can be achieved with near single-cell resolution and demonstrate the feasibility of transcutaneous photoactivation. Future extensions of this approach could enable multicolor, optical reprogramming of cytokine signaling networks and more precise immunotherapies.

## INTRODUCTION

Cytokine signaling is critically important to a variety of physiological processes including cell and tissue development, aging, disease pathogenesis, and the mounting of effective innate or adaptive immune responses.^1-3^ In addition to serving as signal mediators, these proteins can also act as potent therapies with more than 18 cytokine products currently FDA-approved for the treatment of diseases including chronic hepatitis, multiple sclerosis, rheumatoid arthritis, chronic kidney disease, degenerative disk disease, and multiple types of cancer. While cytokines hold great potential as tools to both study and treat human disease, *in vivo* effects of these proteins are often highly pleiotropic and thus difficult to understand and challenging to control.^4^

One mechanism by which cytokines with diverse effects can transmit tissue- and cell-specific information is via expression in an inactive, or *latent*, form in which the protein is sterically shielded by another peptide or protein binding partner. Transforming growth factor-β1 (TGF-β1), for example, has been recently shown to non-covalently associate with latency associated peptide (LAP) which de-shields from TGF-β1 in response to traction forces caused by the binding of αVβ6 or αVβ8 integrins with either cell membrane-bound GARP (glycoprotein A repetitions predominant) or extracellular matrix-bound LTBP-1 (latent transforming growth factor beta-binding protein 1).^5^ This stimuli-responsive uncaging of the protein can lead to remarkable specificity: interaction with migratory dendritic cells has been shown to present TGF-β1 to naïve CD8^+^ T cells, preconditioning them for tissue-resident memory fate.^6^ Similarly, interaction with regulatory T cells (Tregs)^7^ and microglia^8^ has been found to de-shield TGF-β1, thus initiating anti-inflammatory signaling cascades in these cells, as well as other nearby cell types.

Inspired by the ability of reversible shielding to impart cell- and tissue-specificity to cytokines expressed in a latent form, we hypothesized that chemical modification with synthetic macromolecules could impart similar or improved specificity to other cytokines not expressed in a latent state. Photo-responsive linker technologies present a potential, synthetic alternative to latency binding proteins, providing spatiotemporal control over cytokine activation and, additionally, orthogonality to existing shielding/de-shielding pairs. Historically, photo-labile linkers have been utilized to reversibly immobilize peptides and oligonucleotides onto purification resins; however, more recently, this approach has been adapted in order to reversibly cage small molecules, peptides, and nucleic acids.^9,10^ For example, caged neurotransmitters have been used to study memory formation in the brain,^11^ caged peptides complexed with MHC have been exploited to study structural reorganization at the immune synapse,^12^ and caged sgRNA has be utilized to spatially constrain gene editing by CRISPR/Cas9.^13^ Extension of this strategy to immune signaling proteins – which can be hundreds of times larger – thus represents a significant and unaddressed challenge.

Here, we describe a strategy whereby cytokines are chemically modified with photo-labile polymers that mimic the induction of protein latency while attached, then de-shield to recover protein activity in response to monochromatic light exposure. This approach enables both the magnitude and duration of cytokine signals to be tuned on-demand, with high spatial resolution, and can be rapidly adapted to a range of additional cytokine or chemokine proteins. Future extensions of this approach could enable optical reprogramming of cytokine signaling networks and could lead to new immunotherapies that are more tissue-specific and patient-personalized.

## RESULTS

### Artificial Cytokine Latency via Photo-labile Polymer Modification

To assess the feasibility of polymer-induced cytokine latency and subsequent light-induced activation (**Fig 1a-c**), we first examined the cleavage efficiency of two distinct polyethylene glycol (PEG) polymers modified to exhibit fluorescence dequenching following the cleavage of *o*-nitrobenzyl linker derivatives when exposed with blue LED light (**Fig S1a,b**). In phosphate buffer, these polymer cages exhibited fast cleavage kinetics (*k’*∼0.028-0.12 min^-1^) that was both power-dependent and highly wavelength-discriminant (**Fig 1d, S1c-e**). Based on these results, we devised a traceless^14^ chemical modification strategy which appended high molecular weight PEG (20 kDa) to cytokine lysine residues by way of a dialkoxy-substituted 2-nitrobenzyl linker (**Fig S2**), selected due to its red-shifted absorption (**Fig S3**) and enhanced water solubility compared with *o*-nitrobenzyl cages.

**Figure 1.**
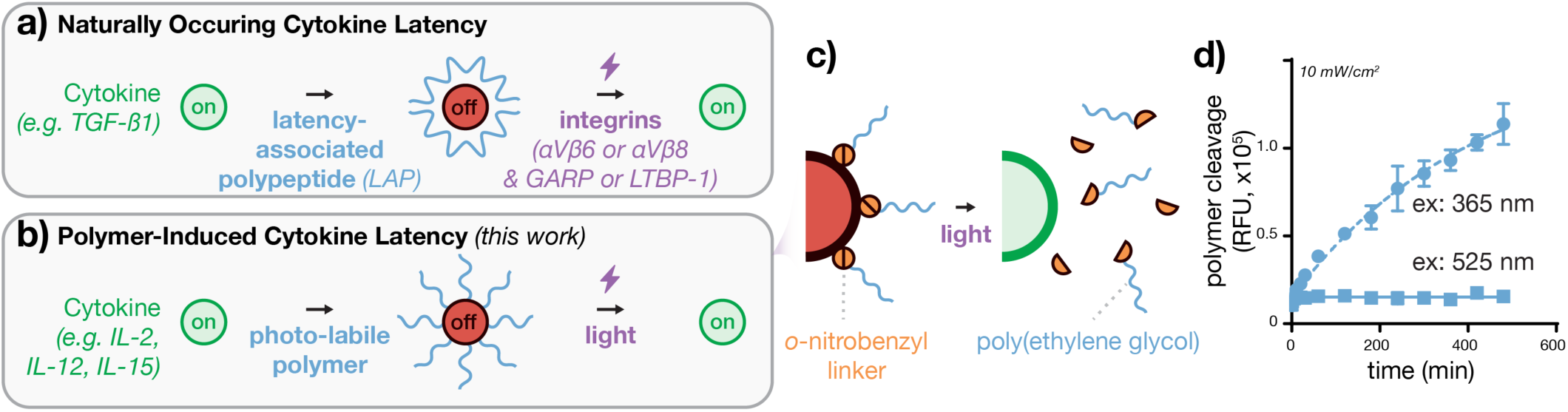
Bioinspired cytokine latency via photo-labile polymer modification. **a)** Transforming growth factor-β1 (TGF-β1) transmits tissue- and cell-specific cytokine signals via association of with latency associated peptide (LAP) which sterically shields and later disassociates from TGF-β1 in response to traction forces caused by the binding of αVβ6 or αVβ8 integrins with either cell membrane-bound GARP or extracellular matrix-bound LTBP-1. **b)** Strategy for the induction of reversible latency in cytokines with pleiotropic effects via modificaiton with end-modified, photo-labile polymers. **c)** Illustration of the traceless modification strategy used here whereby 20 kDa poly(ethylene glycol) polymer chains are appended to cytokine lysine residues via *o*-nitrobenzyl groups which **(d)** are rapidly degraded by blue, but not green, LED light as measured by cleavage-induced fluorescence de-quenching. Data in (c) represent mean±SD of 3 technical replicates.

To demonstrate polymer-induced cytokine latency, we selected recombinant human IL-2 as a candidate for photo-labile polymer modification due to its lack of a known latency binding partner and its well-described pleiotropic effects *in vivo*, for example its simultaneous immunostimulatory effects exerted via cytotoxic T cells and immunosuppressive effects exerted through regulatory T cells. We also selected IL-2 as the protein has demonstrated clinical benefit in patients with melanoma, renal cell cancer, and neuroblastoma. These benefits are greatly limited by the small size and rapid excretion of IL-2 which necessitates continuous or frequent high-dosing and thus toxicity and complex treatment management.^15,16^ *We hypothesized that polymer-induced IL-2 latency could be used to both control IL-2 signaling in vitro and in vivo, as well as improve the safety or therapeutic potential of this and related cytokines via prolonged circulation*.

Following modification of IL-2, we observed a stepwise increase in size upon both linker and polymer conjugation as measured by both polyacrylamide gel electrophoresis and dynamic light scattering (**Fig 2a**). Latent IL-2 was approximately three-fold larger than the wild type protein in overall diameter, thus well above the lower size limit for renal clearance in humans. We further examined the binding affinity of latent IL-2 with two of its cognate receptor subunits, IL-2Rα (CD25) and IL-2Rβ (CD122) via biolayer interferometry (**Fig 2b**). Strikingly, binding affinity of IL-2 towards IL-2Rα was decreased approximately 53-fold following latency-induction, whereas that towards IL-2Rβ was unaltered. This serendipitous result suggests that, relative to wild type IL-2, latent IL-2 maintains biased activity towards CD8^+^ T cells that express IL-2Rβγ and dampened activity towards immunosuppressive Tregs that constitutively express IL-2Rαβγ.^17^ Given that off-target activity towards Tregs is believed to contribute, in part, to the failure of IL-2 therapy in patients,^18^ these findings warrant future investigation in mouse models of cancer and other diseases reliant on T cell immune evasion.

**Figure 2.**
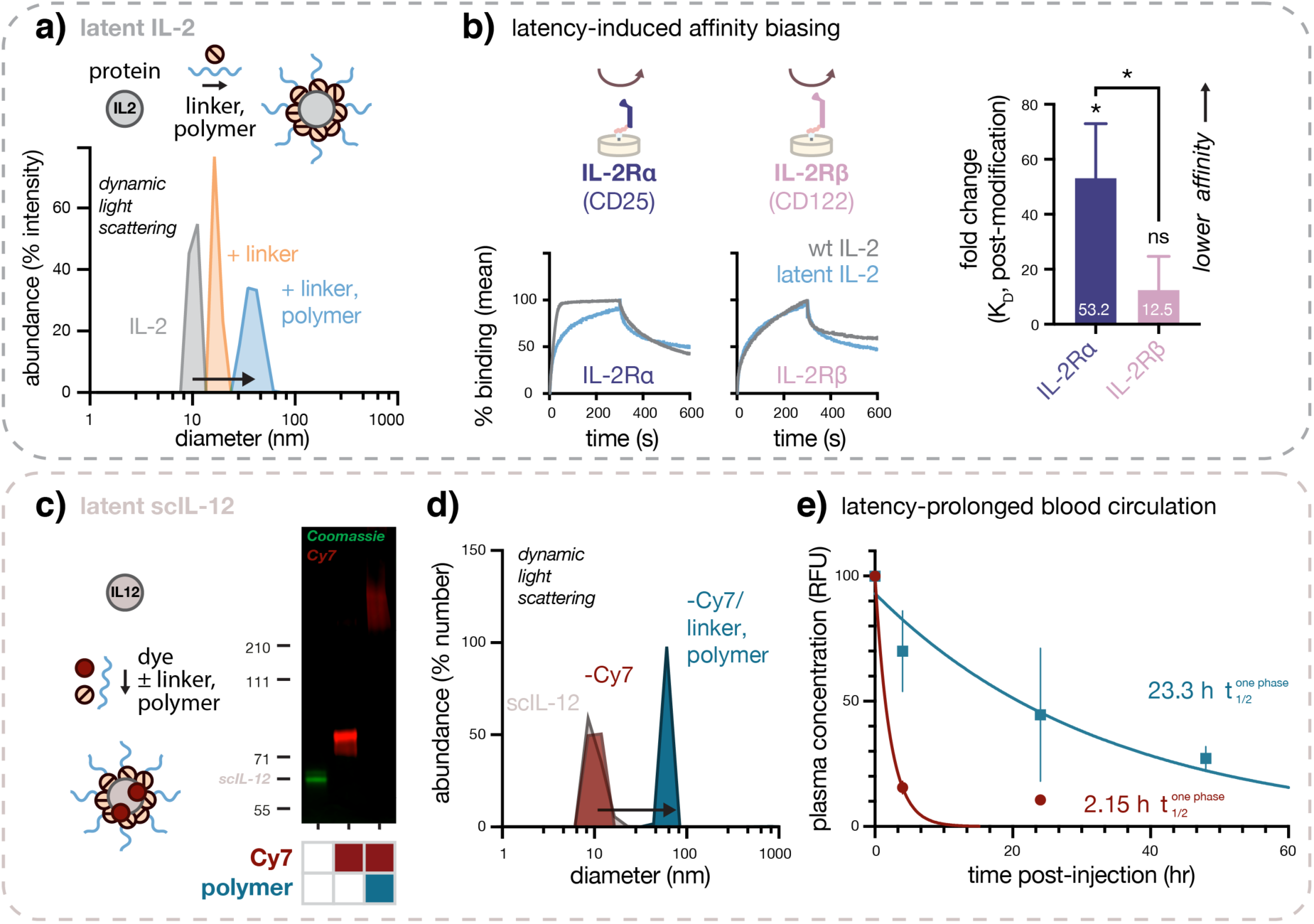
Polymer-induced latency augments cytokine size, biases receptor subunit binding affinity, and prolongs *in vivo* circulation. **a)** Step-wise increase in IL-2 hydrodynamic size upon linker and polymer conjugation as measured by dynamic light scattering. **b)** Sensorgrams depicting binding kinetics for IL-2 or latent IL-2 association/dissociation with IL-2Rα (CD25) or IL-2Rβ (CD122). (Inset) fold-change in post-modification binding affinity. **c)** Electrophoretic mobility shift demonstrating Cy7 dye- and polymer- dependent modification of scIL-12 as measured by polyacrylamide gel electrophoresis. **d)** Increase in scIL-12 hydrodynamic size following Cy7 conjugation with or without linker/polymer modification as measured by dynamic light scattering. **e)** Plasma pharmacokinetics of Cy7-labeled scIL-12 modified with or without linker/polymer following intravenous injection into (C57BL/6 mice) illustrating prolonged circulation following photo-labile polymer modification. Data in (b) represent mean±SD of 3 technical replicates. Data (e) represent mean±SEM of 2-3 biological replicates. p < .05(*), p < .01(**), p < .001(***), p < .0001(****).

Having demonstrated that polymer-induced latency can modulate cognate receptor binding affinity, we characterized the effect of polymer-induced latency on *in vivo* cytokine circulation using IL-12, another recombinant cytokine under clinical investigation which similarly suffers from rapid clearance and systemic, off-target toxic effects.^19^ As therapeutic cytokines are generally quite small (ca. 12-70 kDa), polymer modification – frequently, with PEG – is often used to decrease renal clearance, thus prolonging circulation and augmenting tissue exposure with drug (e.g. Pegfilgrastim, Peginterferon, etc).^20^ To monitor cytokine circulation *in vivo*, we dye-labeled a novel single-chain variant of the cytokine, scIL-12, both with and without photo-labile polymer modification (20 kDa PEG, **Fig 3c**). Cy7-labeling only nominally increased scIL-12 size as measured by polyacrylamide gel electrophoresis and dynamic light scattering, whereas combined dye and photo-labile polymer modification increased hydrodynamic size to 44 nm (**Fig 3d**), well above the renal clearance size threshold in humans and rodents. We then monitored the circulation of latent scIL-12 following tail vein injection in C57BL/6 mice, finding that the latent cytokine experienced 16-fold increase in circulation half-life following polymer-modification (**Fig 3e**). These results suggest that the prolonged circulation of scIL-12 may (i) obviate the need for frequent, high dosing and (ii) improve tissue accumulation of the drug in therapeutic settings.

**Figure 3.**
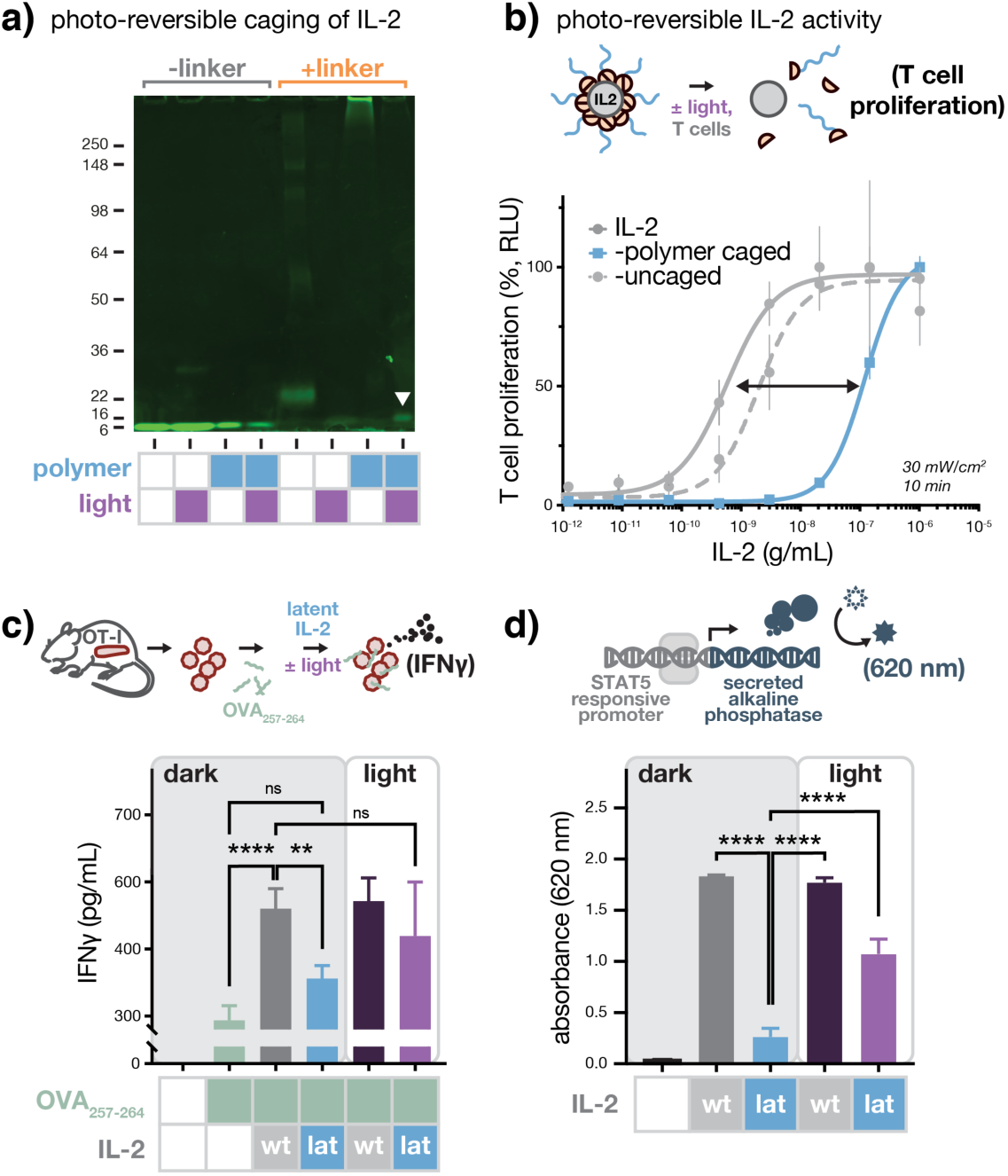
Photo-exposure of latent IL-2 restores protein size and functional activity. **a)** Electrophoretic mobility shift demonstrating linker- and polymer- dependent modification of IL-2, as well as light-dependent restoration (arrowhead) of wild-type protein mobility as measured by polyacrylamide gel electrophoresis. **b)** Polymer-dependent repression and light-induced restoration of IL-2 activity as measured by CTLL-2 T cell proliferation (24 h). **c)** Effect of latent IL-2 and uncaged IL-2 (1000 IU/mL molar equivalents) on OVA_257–264_ antigen-specific T cell activation as measured *ex vivo* by ELISA of OT-I splenocyte-secreted interferon gamma (IFN*γ*). Data represent (b) mean±SD of 3 technical replicates and (c) mean±SD of of 6 technical replicates. p < .01(**),p < .0001(****).

### Photo-Activation of Latent Cytokines

After characterizing the effects of polymer-induced cytokine latency, we next examined the recovery of functional protein activity using CTLL-2 T cells that depend on both IL-2 and IL-2Rα for growth.^21^ Following latency-induction, we observed an approximate 10^3^-fold drop in rhIL-2 activity as measured by CTLL-2 T cell proliferation (24 h); however, subsequent LED irradiation to uncage IL-2 near fully restored both native molecular weight and capacity for induced T cell proliferation (**Fig 3a,b**). *In vivo*, as little as a 10-fold change in rhIL-2 activity is necessary in order to functionally modulate fate decisions in T cells that lead to either memory or effector fate,^22^ thus the near three logs of dynamic range observed here *in vitro* suggest that latent cytokines may be used to control T cell biology *ex vivo* or modulate therapeutic activity of the recombinant protein. To further demonstrate the feasibility of this approach, we also examined latency-induction with rhIL-15, finding that small molecule linker addition, alone, was sufficient to achieve 5- to 20-fold modulation of CTLL-2 T cell dose-dependent proliferation (24 h, **Fig S4**).

To further explore the therapeutic potential of latent IL-2, we investigated its ability to promote antigen-specific immunity using OT-I T cell receptor transgenic mice which generate clonal CD8^+^ T cells specific to SIINFEKL, an octameric peptide from ovalbumin (OVA_257–264_).^23^ We pulsed OT-I splenocytes with OVA_257-264_ and treated with either wild type or latent IL-2, with or without LED irradiation, and measured IFN*γ* as an indication of the extent of T cell activation. While latent IL-2 had no significant effect on antigen-specific T cell activation, that from the light-uncaged protein was comparable and statistically indistinguishable from wild type IL-2 (**Fig 3c**).

To ascertain whether the activity of latent IL-2 on OT-I T cells was, like the wild type protein, JAK/STAT pathway-dependent, we examined its effect on HEK293 cells engineered to express all three subunits of human IL-2R as well as JAK3 and STAT5. In response to STAT5 activation, these cells secrete alkaline phosphatase which can be spectrophotometrically detected using a chromogenic substrate. The trends in reporter cell response observed in these studies closely match those observed in activated OT-I T cells. Latent IL-2 induced near baseline levels of STAT5 transcriptional activity, while that from the light-uncaged protein was comparable to that from wild type IL-2 (**Fig 3d**). Together, these data support that LED irradiation can de-repress IL-2 latency, promote antigen-specific immunity, and re-activate JAK/STAT pathway signaling.

### Feasibility of In Situ Cytokine De-Repression

Having shown that cytokine latency can be used to temporally control immune cell signaling, we sought to characterize the spatial resolution with which cytokine activity could be constrained. We prepared a latent analog of IL-2 using bovine serum albumin modified with a photo-labile PEG containing a biotin tag at its distal end (**Fig 4a**). Following immobilization onto glass slides, irradiation through a custom photolithographic mask, and streptavidin-FITC staining, we observed spatially constrained protein uncaging with resolution at or below the typical dimensions of single human immune cells (<17 μm, **Fig 4b,c**). These results suggest that cytokine activity can be de-repressed with high spatial *and* temporal control using this synthetic modification approach.

**Figure 4.**
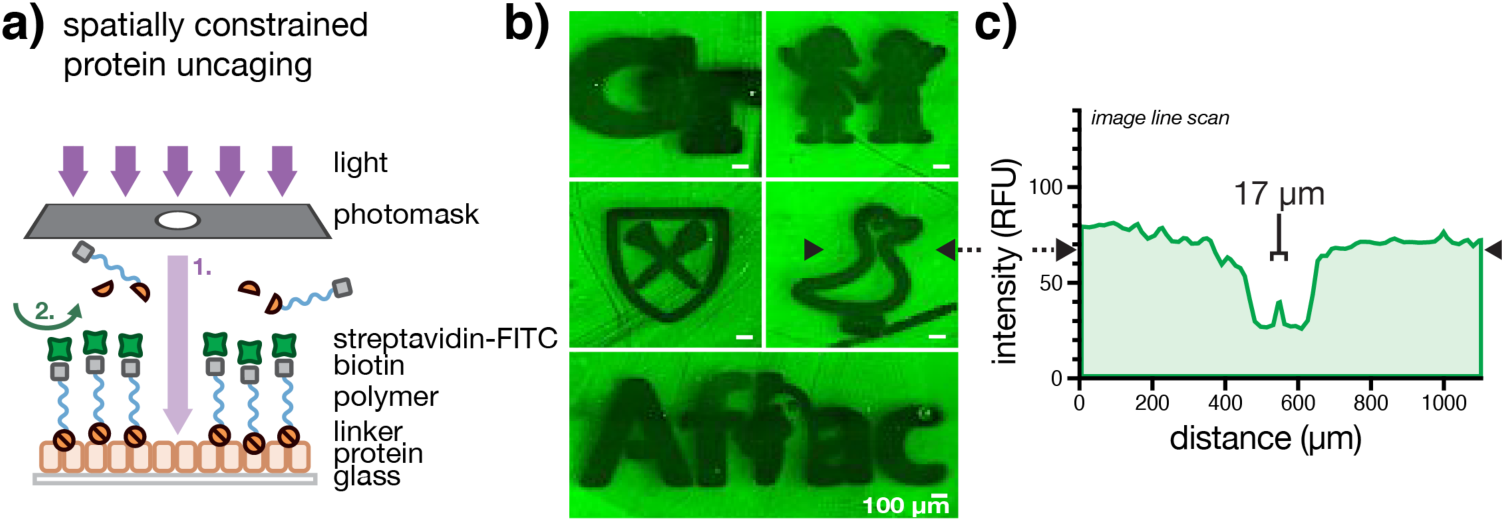
Light-induced uncaging enables precise, local control of protein activity. **a)** Illustration of experiments to visualize protein latency and photo-induced de-repression. **b)** Fluorescence micrographs and **(c)** corresponding image line scan image indicating regions of latent (green) and uncaged (black) protein as measured by epifluorescence microscopy.

To explore the effects of tissue light attenuation on latent IL-2 activation, we fabricated a series of silicone-based phantoms that mimic light transmission through human dermis, epidermis, and hypodermis at wavelengths specific to the polymer photocages described here (**Fig 5a**). Using these models, we examined the stability of 5 kDa PEG photocages under prolonged, aqueous storage conditions and under conditions simulating ambient indoor light exposure of superficial veins (1 mm hypodermis depth^24^). We observed high stability of aqueous solutions in cold storage with <10% total uncaging of polymer linkages after 90 days (**Fig 5b**). Given that many clinical products have post-reconstitution shelf-lives of just hours to days, the storage durations observed here appear sufficient for large-scale *in vivo* testing. We also observed comparably low levels of polymer cleavage under conditions mimicking non-deliberate light exposure of superficial veins over 6 weeks (<4%). These latter data suggest that polymer-induced latency may be maintained *in vivo* over time scales necessary for light-constrained cytokine de-repression.

**Figure 5.**
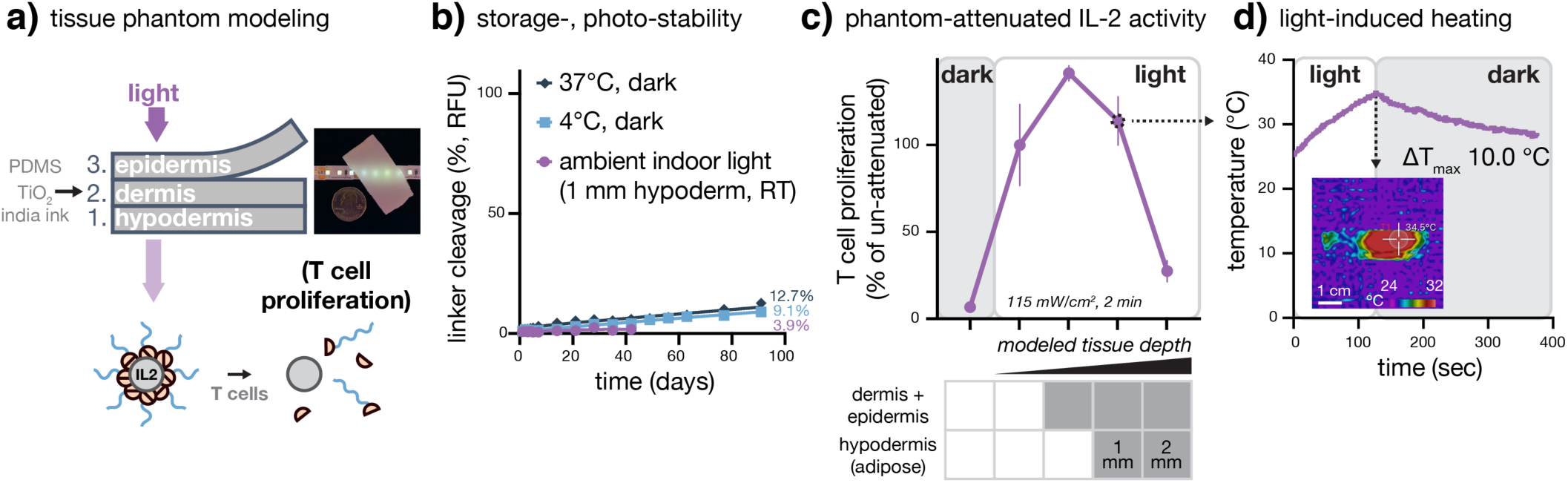
Latent IL-2 is stable and feasibly photo-activated through tissue models. **a)** Illustration of multilayered tissue phantom construction. **b)** Stability of 5 kDa poly(ethylene glycol) polymer photocages under prolonged, aqueous storage and under conditions simulating ambient, indoor light exposure of superficial veins (1 mm depth) as measured by cleavage-induced fluorescence dequenching. **c)** Effection of tissue phantom light attention on latent IL-2 activity as measred by CTLL-2 T cell proliferation (24 h). **d)** Superficial heating of multilayered tissue phantoms of the indicated thickness as measured by forward-looking infrared imaging. Data in (c) represent mean±SD of 3 biological replicates. p < .05(*), p < .01(**), p < .001(***), p < .0001(****).

To model the feasibility of light-induced cytokine uncaging *in vivo*, we examined the activity of latent IL-2 following photo-exposure through tissue phantoms modeling human skin and subcutaneous tissue. We observed near full recovery of IL-2 activity at depths corresponding to 1 mm beneath the dermis as measured by CTLL-2 T cell proliferation (**Fig 5c**). These findings are significant as such depths, in many cases, correspond to the minimal light attenuation experienced at human superficial veins as well as within transcutaneous or some transepithelial tumors.^24^ Moreover, as the light irradiance required for activation through tissue phantoms induced only a small temperature increase (ΔT_max_ 10.0 °C, **Fig 5d**), heat pain responses *in vivo* are expected to be mild or imperceptible.^25,26^

## DISCUSSION

Inspired by the ability of latency binding to impart specificity to otherwise pleiotropic immune signaling proteins, here we describe a strategy whereby chemical modification with light-sensitive polymers can be used to control the magnitude, duration, and location of cytokine signals in response to simple LED light exposure. In this study, we found that modification of IL-2 and IL-15 with photo-labile small molecules or polymers could modulate their activity on T cells as much as two to three orders of magnitude. This ability to control the magnitude, and correspondingly the duration, of IL-2 signal is significant as (i) both strong and sustained IL-2 signaling is necessary for the induction of CD8^+^ effector or memory T cell fate, and (ii) as little as a ten-fold change in local cytokine concentration can bias this tradeoff in short-term and long-term immunity.^22,27^ Strategies such as those described here may, in future work, enable the optical reprogramming of fate decisions in T cells confer short-term effector function at the expense of long-term memory function.

In this work, we also show that polymer-induced latency blunts corresponding JAK/STAT pathway signaling and CD8^+^ T cell activation *ex vivo*, and that just brief LED light exposure can be used to de-repress these effects. We envision that such high spatial and temporal control of cytokine signaling can be used modulate T cell priming/expansion directly at sites of disease or at associated secondary lymphoid organs. Such strategies may also extend to chemokines which can serve to further orchestrate effective adaptive immune responses against pathogens or tumors. Here, we achieved a minimum spatial resolution of photoactivation approaching that of a single immune cell – without the use of focusing optics – and although light scattering and diffusion would limit such dimensions *in vivo*, we anticipate strong feasibility to constrain cytokine activation to mm-scale diseased tissues and lymph nodes in future work. This supposition is also supported by tissue phantom studies performed here, showing efficient light-induced de-repression at subdermal depths of as high as 1 mm, sufficient in many instances for activation within human superficial veins as well as within transcutaneous or some transepithelial disease sites.^24^ Others have also demonstrated that structurally related photocages appended to solid implants can be transcutaneously photoactivated in mice,^28,29^ thus we are optimistic regarding future *in vivo* feasibility.

Serendipitously, we also found that polymer-induced latency, alone, biased the affinity of latent IL-2 towards IL-2Rβ (CD122) and away from IL-2Rα (CD25). As CD8^+^ T cells express IL-2Rβγ and immunosuppressive Tregs constitutively express IL-2Rαβγ, these findings suggest that the latent cytokine may improve CD8^+^/Treg ratios which are prognostically favorable in many cancers^30^ and correlate with clinical responses to immune checkpoint blockade therapy in patients.^31^ While other mechanisms of cytokine receptor subunit-biasing based on mutagenesis,^32,33^ antibody complexation,^34^ and *de novo* protein design^35^ have been reported, here, we hypothesize that atypically high density of solvent-accessible lysine resides at the IL-2/IL-2Rα interface preferentially induce steric hindrance with the receptor subunit via appended polymer chains.^17^

In addition to demonstrating rapid and efficient cytokine photo-activation, we also found that polymer-photocages used here were highly stable under conditions simulating both aqueous storage and venous ambient light exposure. Compared with other promising and more rapidly hydrolyzable PEG/IL-2 conjugates (*e*.*g*. NKTR-214^36^), these findings are encouraging and could lead to future integration with wearable or implanted light-delivery devices^37-41^ which modulate drug activation. Consistent with other PEGylated cytokines,^36^ we found that bioinspired, polymer-induced latency was able to prolong scIL-12 plasma circulation approximately 16-fold, potentially precluding the need for frequent, high dosing and improving tissue drug exposure in treatment settings.

While these studies are the first, to our knowledge, to demonstrate reversible, optical control of cytokines, they build upon many prior advances.^9-13,42-46^ Deiters and coworkers previously demonstrated reversible photocaging strategies for enzymes based on nitrobenzyl linkages,^47^ while Esser-Kahn and coworkers have demonstrated related approaches to photo-activate smaller lipopeptide Toll-like receptor (TLR) agonists.^48,49^ Likewise, Garcia,^50^ Hubbell,^51^ and Wittrupp^52^ have recently developed affinity-targeted cytokine fusions with high cell- or tissue-specific activity. Combining elements of these approaches, here we describe a bioinspired strategy for polymer-induced cytokine latency and photo-induced reactivation. The work presented here provides proof-of-concept that cytokine activity can be precisely regulated using light, findings that may be further improved upon through the use of additional photo-labile linkers which absorb light at wavelengths with higher tissue penetrance.^9,53-55^

In summary, we describe a generalizable strategy for light-induced cytokine de-repression that can be used to spatially and temporally control the activity of otherwise pleiotropic immune signaling molecules. As research tools, the technologies described here hold great potential to improve our ability to understand and manipulate basic immune biology, and to optically reprogram immune responses *ex vivo* and *in vitro*. As therapies, they could also serve as long-acting pro-drugs or tissue-selective immune modulators both alone or in combination with other immunotherapies.

## METHODS

### Materials and supplies

Unless otherwise specified, reagents were used as received without further purification. Recombinant human IL-2 (200-02, Peprotech), recombinant human IL-15 (570308, Biolegend), recombinant mouse scIL-12 (130-096, Miltenyi Biotech). Sulfo-Cyanine7 NHS ester (Lumiprobe), DBCO-NHS ester (1160, Click Chemistry Tools), poly(ethylene glycol) methyl ether azide (20 kDa, Nanocs). Polyacrylamide gels (Bio-Rad, 4-16 wt%). IFNγ ELISA (DY485, R&D Systems).

### Photo-induced polymer cleavage

Detailed characterization of light-induced polymer cleavage was monitored via fluorescence de-quenching of 5FAM- and CPQ2-modified polyethylene glycol (5 kDa). Polymers containing the photo-labile linkers shown in Fig S1b were obtained from CPC, Inc using linker reagents obtained from Advanced Chemtech. Modified polymers (azido-G-K(CPQ2)-NB/DMNB-PEO4-G-K(5FAM)-GC-PEG5k) were >96% pure as measured by RP-HPLC. Samples were dissolved to 500 nM in ultrapure water and irradiated within quartz cuvettes using collimated, light-emitting diodes (Solis, Thorlabs). Fluorescence de-quenching was measured on a Spectramax Id3 plate reader (Molecular Devices). Cleavage kinetics were fit to a one-phase decay using Graphpad Prism software. Storage and stability measurements were similarly obtained from solution aliquots maintained in on a laboratory benchtop, heated bead bath, or laboratory refrigerator. Samples were covered with aluminum plate film or tissue phantoms and exposed to fluorescent, overhead office lights as indicated.

### Polymer-induced latency

Recombinant cytokines were sequentially modified with photo-labile linkers and polymers via carbodiimide coupling and Cu-free click chemistry, respectively. Briefly, cytokines were reacted with a commercial 2-nitrobenzyl linker displaying both NHS ester and DBCO substituents (1160-10, Click Chemistry Tools) followed by addition of poly(ethylene glycol) methyl ether azide. Reaction conditions are described as molar equivalents relative to total lysine residues or total DBCO groups.

Recombinant scIL-2 was modified via dilution in 150 mM sodium phosphate buffer (pH 8.5) containing 0.5 mM SDS and addition of 3 eq. of photo-labile linker to a final DMSO concentration of 5% v/v. 10 eq. of poly(ethylene glycol) methyl ether azide (20 kDa) dissolved in PBS (pH 7.4) was then added and allowed to react. All cytokine modification steps were allowed to proceed overnight at 4 °C with rotary agitation (800 rpm). In some experiments, excess PEG was removed via DBCO-agarose beads (Click Chemistry Tools).

Recombinant IL-12 was modified via dilution in 150 mM sodium phosphate buffer (pH 8.5) containing 0.5 mM SDS and addition of (i) 1 eq. of NHS-sulfoCy7 to 5% v/v DMSO or (ii) addition of 1eq. of NHS-sulfoCy7 and 5 eq. of photo-labile linker to 5% v/v DMSO. 5 eq. of poly(ethylene glycol) methyl ether azide (20 kDa) dissolved in PBS (pH 7.4) was then added and allowed to react. Excess dye was removed via desalting column (7k MWCO, Pierce).

Recombinant IL-15 was modified via dilution in buffer containing 10 mM NaH_2_PO_4_/150 mM NaCl and addition of 20 eq. of photo-labile linker to 5% v/v DMSO.

### Protein characterization

Cytokine hydrodynamic size was measured by dynamic light scattering (Wyatt Dynamo Plate Reader III). Electrophoretic mobility was measured via polyacrylamide gel electrophoresis under reducing conditions. Protein bands were visualized with Coomassie G250 stain (Bio-Rad) and imaged using a Licor CLx gel imager.

### Cytokine-induced proliferation

Murine CTLL-2 T cells (ATCC) were maintained in RPMI 1640 with high glucose, L-glutamine, and HEPES and supplemented with 10% heat inactivated FBS, 10% rat T-STIM (Corning), 2 mM L-glutamine, and 1 mM sodium pyruvate. To examine cytokine activity, cells were washed in assay media (maintenance media without T-STIM) and plated in assay media at 1.5×10^3^ cells per well within a 96-well plate. Wells were then treated for 24 h with equimolar amounts of wild type or latent protein that was LED-exposed for the indicated time/irradiance. Cell proliferation was measured using CellTiter-Glo 2.0 reagent (Promega) following the protocol provided by the manufacturer. Cell lines were routinely screened for mycoplasma (MycoAlert Plus, Lonza).

### Antigen-specific T cell activation

OT-I mouse (6-8 wk) splenocytes were isolated using Ficol-Paque following mechanical homogenization and PBS washing of excised spleen tissues. Spleenocytes were resuspended in RPMI 1640 (ATCC) containing 10% FBS, 100 U/mL penicillin G and streptomycin. For stimulation, cells were incubated with 10 nM chicken egg ovalbumin peptide 257-264 (SIINFEKL, Invivogen) with or without 1000 IU/mL IL-2 (or equimolar amounts modified protein) and plated at 2×10^6^ cells/well within a 96 well plate. After 24 hours, cell culture supernatant was harvested and tested for IFNγ secretion via ELISA (R&D Systems). Optical density at 450 nm was measured using a Spectramax Id3 plate reader (Molecular Devices) and results were compared with standard curves. These studies were approved by Emory University’s Institutional Animal Care and Use Committee.

### JAK/STAT pathway activation

STAT5:SEAP reporter cells (HKB-il2, Invivogen) were maintained in DMEM supplemented with 4.5 g/L glucose, 2-4 mM L-glutamine, 10 %v/v heat-inactivated FBS, 100 U/ml penicillin, 100 μg/ml streptomycin, 100 μg/ml Normocin, and HEK blue selection media (Invivogen). For assays, 5×10^4^ cells were plated in complete growth media, without selection antibiotics, within 96-well plates and treated with equimolar amounts of wt or latent IL-2 for 48 h. Then, 20 *µ*L of cell culture supernatant was then withdrawn and analyzed for alkaline phosphatase content via change in Quanti-Blue (Invivogen, rep-qbs) absorbance at 620 nm (Spectramax id3 plate reader).

### Binding affinity

Cytokine and cognate receptor binding kinetics were measured via biolayer interferometry using an Octet RED384 system (ForteBio). Nickel nitrilotriacetic acid sensors (Ni-NTA, ForteBio) were equilibrated in PBS and coated with polyhistidine-tagged receptor proteins (SinoBiological) for 5 minutes (1.5 μg/mL IL-2Rα, 3.3 μg/mL IL-2Rβ). Association kinetics were measured over 5 min at 2.0 μM IL-2, followed by dissociation in PBS over 5 min. Measurements were repeated in three independent experiments and fit using Data Analysis 8.0 (ForteBio) using a 1:1 kinetic model.

### Pharmacokinetics

C57BL/6 mice (female, 7 wk) were injected via tail vein with 2 ug scIL-12 protein conjugated with Cy7 alone or Cy7 with photo-labile poly(ethylene glycol). Plasma was collected from 100 μL of blood obtained via submandibular bleed into heparinized tubes (BD Microtainer). Plasma fluorescence was integrated using ImageJ software after polyacrylamide gel electrophoresis. Mice with peak plasma fluorescence <1.2x above baseline were excluded. These studies were approved by Emory University’s Institutional Animal Care and Use Committee.

### Tissue phantoms

Polydimethylsiloxane (PDMS) tissue phantoms were prepared as described previously.^56-58^ Briefly, Sylgard 184 elastomer and curing agent (Dow Corning) was doped with titanium dioxide (0.3-1.0 μm rutile, Atlantic Equipment Engineers) and india ink (Higgins 44201) to concentrations which approximate attenuated light transmission through human tissue (36, 20, and 42% transmission of 365 nm light through epidermis, dermis, and 2 mm hypodermis tissue, respectively).^59^ Solutions were degassed and sequentially cast into rectangular silicone molds prior to measurement of LED light transmission using a thermal power sensor (ThorLabs). Mean light transmission through individual phantom layers was measured as: 33.3 %T (epidermis), 28.1 %T (dermis), 39.3 %T (hypodermis). Photo-induced heating measurements were collected using a FLIR ONE Pro thermal imaging camera and analyzed using Vernier Thermal Analysis Plus software.

### Statistical analysis

All p-values were calculated using either two-way ANOVA with Tukey post-hoc correction or two-way T test, depending on sample number, using Graphpad Prism software unless otherwise specified.

### Photo-patterned protein activation

Silicone cell isolators (Electron Microscopy Sciences) were adhered to aldehyde-functionalized glass slide (Nanocs) and coated overnight in 2 mg/mL BSA (VWR) at RT. Coated wells were then washed 2x with 0.2% SDS and 2x with deionized water. Protein was covalently bound to the slide surface via addition of freshly prepared 2.5 mg/mL NaBH_4_ (Sigma) dissolved in 25 %v/v ethanol in PBS at RT for 5 min. Wells were again washed 3x with 0.2% SDS and 3x with deionized water. DBCO-sulfo-NHS ester (Sigma) in PBS was added at 5 eq. (relative to lysine residues) for 4 hr at RT, then wells were again washed 3x with deionized water. Azide-PC-Biotin (Click Chemistry Tools) dissolved in DMSO was added to the wells at 1 eq. (relative to DBCO groups) and incubated overnight at RT. Wells were washed 3x with deionized water and filled with 50% glycerol prior to UV exposure of through custom, chrome-patterned quartz photomasks (Front Range Photomask) for 30 min with a 365 nm LED (Thor Labs) at 30 mW/cm^2^. Wells were washed 6x with deionized water, 3x with 0.2% SDS, and again 3x with deionized water. Streptavidin-FITC (Southern Biotech) in PBST (0.1% Tween 20) was incubated in the wells for 30 min at RT, then wells were washed 3x with PBST and 3x with deionized water. Cell isolators were removed, ProLong Diamond Antifade mounting media was added (Life technologies), and the slide was coverslipped. The slide was visualized using a widefield microscope with GFP filter settings (469/525 nm, Biotek Lionheart FX) and images were processed with Gen5 and ImageJ software.

## ACKNOWLEDGEMENTS

This work was supported in part by the American Cancer Society (#IRG-17-181-04), the Winship Cancer Institute, the National Institute of Health Research Training Program in Immunoengineering (T32EB021962), the AAI Careers in Immunology Fellowship Program, the Coulter Department of Biomedical Engineering, and the Aflac Cancer and Blood Disorders Center of Children’s Healthcare of Atlanta. We are also grateful for assistance from the Children’s Healthcare of Atlanta and Emory University’s Pediatric Integrated Cellular Imaging Core and Pediatric General Equipment & Specimen Processing Core, the Robert P. Apkarian Integrated Electron Microscopy Core, and the Emory Chemical Biology Discovery Center. The content here is solely the responsibility of the authors and does not necessarily represent the official views of the Winship Cancer Institute, Aflac Inc., Children’s Healthcare of Atlanta, or the National Institutes of Health.

## AUTHOR CONTRIBUTIONS

L.A.P., P.D., K.S., G.B.L., C.C.P., and E.C.D. designed research; L.A.P., P.D., C.D., A.C., A.K., A.R., and H.K., K.S., G.B.L., C.C.P, and E.C.D. performed research or analyzed data; and L.A.P., C.C.P., and E.C.D. wrote the manuscript.

