## Supplemental Information for "Optical Control of Cytokine Signaling via Bioinspired, Polymer-Induced Latency"

### SUPPLEMENTARY INFORMATION

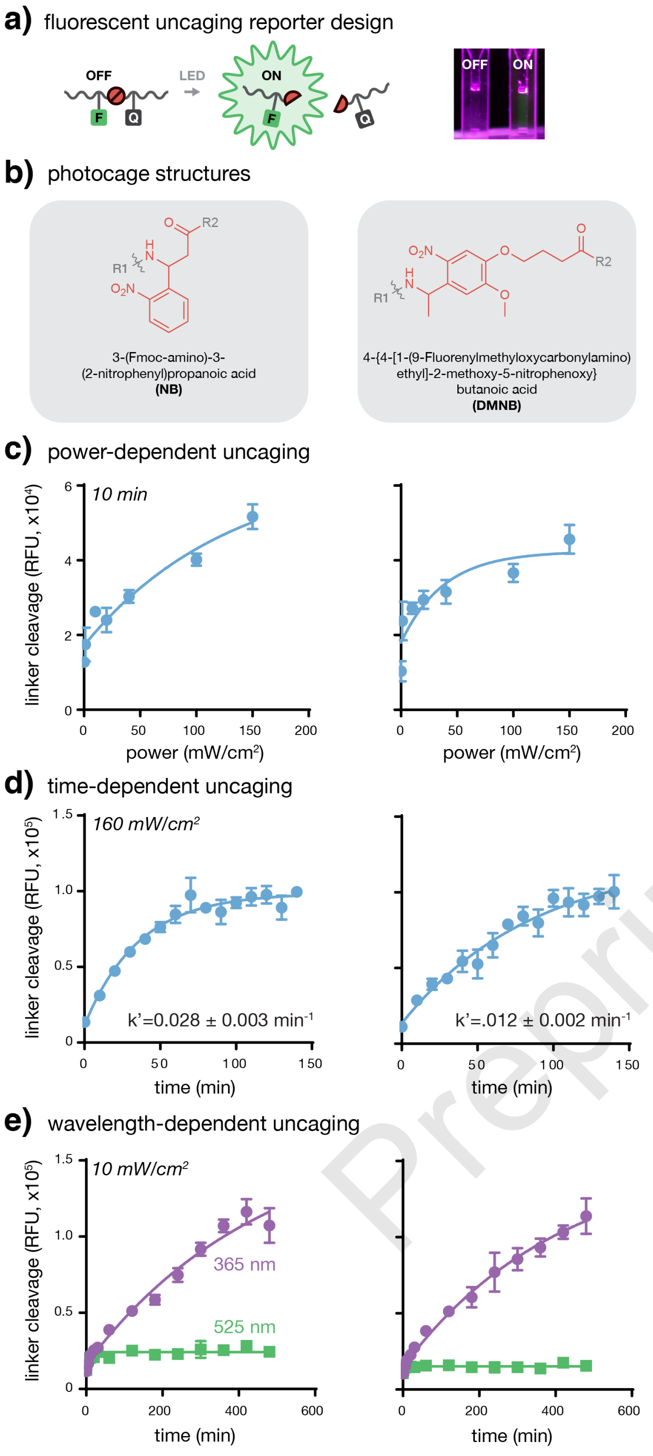


**Figure S1. Light-induced cleavage of two *o*-nitrobenzyl-linked poly(ethylene glycol) polymers is efficient and wavelenth-discriminant. a)** Illustration of cleavage-induced fluorescence de-endenching reporters and (inset) optical images of images of irridiated and non-irridiated samples. **b)** Chemical structures of (left) NB and (right) DMNB photocages. **c)** Power-, **(d)** time-, and **(e)** wavelength-dependence of polymer photocleavage as measured by fluorescence dequenching. Data in (c-e) represent mean±SD of 3 technical replicates.

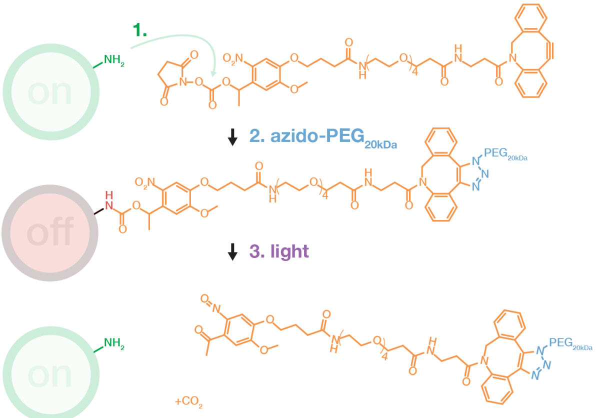


**Figure S2. Photo-labile polymer modification of cytokines.** Traceless modification strategy whereby cytokine lysine residues are modified with NHS-nitrobenzyl-DBCO groups via carbodiimide coupling, followed by conjugation with mPEG-$N_3$ (20 kDa) via copper-free click chemistry.[3]

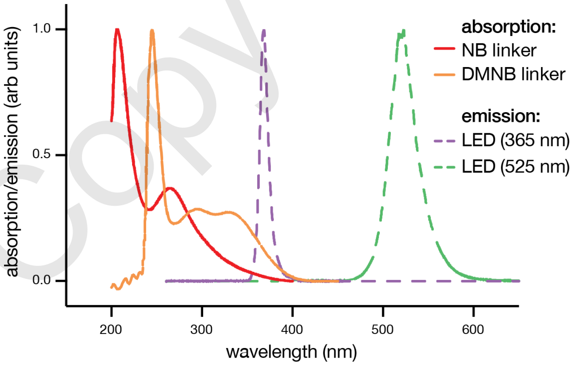


**Figure S3. Optical properties of photosensitive linkers and light sources.** Absorption (solid) and emission (dashed) spectra of photo-labile linkers and light emitting diode (LED) sources as approximated from structural anlogs. Data adapted from Thorlabs (LEDs),[1] *N*-Nmoc-L-glutamate (*e.g.* NB),[2] and *N*-Benzyl-(4-(1-acetoxyethyl)-2-methoxy-5-nitrophenoxy)acetamide (*e.g.* DMNB).[4]

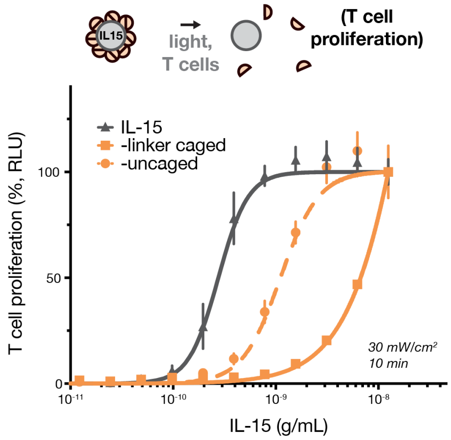


**Figure S4. Photo-exposure of latent IL-15 restores cytokine activity on T cells.** Linker-dependent repression and light-induced restoration of IL-2 activity as measured by CTLL-2 T cell proliferation (24 h). Data represents mean±SD of 3 technical replicates.

Preprint copy


Preprint Copy
